# Rapid profiling of transcription factor-cofactor interaction networks reveals principles of epigenetic regulation

**DOI:** 10.1101/2024.04.05.588333

**Authors:** MM Inge, R Miller, H Hook, D Bray, JL Keenan, R Zhao, TD Gilmore, T Siggers

## Abstract

Transcription factor (TF)-cofactor (COF) interactions define dynamic, cell-specific networks that govern gene expression; however, these networks are understudied due to a lack of methods for high-throughput profiling of DNA-bound TF-COF complexes. Here we describe the <u>Co</u>factor <u>Rec</u>ruitment (CoRec) method for rapid profiling of cell-specific TF-COF complexes. We define a lysine acetyltransferase (KAT)-TF network in resting and stimulated T cells. We find promiscuous recruitment of KATs for many TFs and that 35% of KAT-TF interactions are condition specific. KAT-TF interactions identify NF-κB as a primary regulator of acutely induced H3K27ac. Finally, we find that heterotypic clustering of CBP/P300-recruiting TFs is a strong predictor of total promoter H3K27ac. Our data supports clustering of TF sites that broadly recruit KATs as a mechanism for widespread co-occurring histone acetylation marks. CoRec can be readily applied to different cell systems and provides a powerful approach to define TF-COF networks impacting chromatin state and gene regulation.

## Introduction

Gene expression is coordinated by TF binding to cis-regulatory elements (CREs) throughout the genome.^1,2^ TF function is subsequently carried out by the recruitment of COFs that are not classical DNA-binding TFs^2^ but are recruited to DNA by protein-protein interactions (PPIs) with DNA-bound TFs to modulate transcription. COFs perform diverse functions related to gene regulation, including modifying histones, remodeling chromatin, and interacting with the transcriptional machinery.^1,3–6^ TF-COF interactions can be cell type specific,^7,8^ can change in response to different signals,^9–13^ and can be altered in a range of diseases.^7,14,15^ As such, TF-COF interactions define a dynamic network that governs gene regulation in the cell. Nevertheless, despite their central role in coordinating cellular responses, there is surprisingly limited information about cell-specific TF-COF networks, due primarily to a lack of methods for high-throughput profiling of TF-COF complexes.

Multiple high-throughput methods exist for assaying PPIs; however, each has features that complicate their use for rapid profiling of cell-specific, DNA-bound TF-COF complexes. The yeast two-hybrid assay^16–18^ is widely used to profile pairwise PPIs but is unable to evaluate cell-type-specific interactions as the experiments are performed in yeast. Protein-affixed arrays provide an alternative approach to screen for PPIs, but the need to deposit purified proteins onto a microarray complicates measurement of interactions that require cell-specific, post-translational modifications.^19^ Proximity labeling,^20,21^ affinity purification followed by mass spectrometry (AP/MS),^22–26^ and mammalian two-hybrid^27^ approaches all provide the ability to profile PPIs in a cell-specific context. However, these approaches all require cloning protein tags or enzymes onto target proteins and expressing them uniformly in mammalian cells, which hinders their use for rapid profiling of factors in different cellular contexts. A final complication with these available approaches, with the exception of proximity labeling, is that they do not assay PPIs in the presence of DNA and therefore will miss TF-COF interactions that require TFs to be in a DNA-bound conformation, which can occur for regulatory complexes.^28,29^

In this work, we describe the CoRec approach for the rapid profiling of cell-specific, DNA-bound TF-COF complexes. CoRec builds upon our recent work applying the protein-binding microarray (PBM) technology to study TF-COF complexes present in cell nuclear extracts.^30–32^ We use CoRec to profile cell-specific TF-COF interactions for different classes of COF. In addition, we use CoRec to examine KAT-TF interaction networks in resting and stimulated T cells and determine how these networks relate to mechanisms of epigenetic regulation. We demonstrate that profiling KAT-TF interactions can identify both constitutive and stimulus-dependent TF activators whose DNA binding sites correlate with histone acetylation levels genome-wide. We identify many KAT-TF interactions that have not been previously reported, suggesting that TF interactions with many COFs are understudied. Finally, we identify promiscuous KAT interactions with many TF families and show that heterotypic clusters of binding sites for KAT-recruiting TFs predict differential promoter acetylation.

## Results

### CoRec is an array-based method to profile cell-specific TF-COF interactions

To examine TF-COF networks in different cell types and stimulus conditions, we have developed the CoRec method to profile TF-COF complexes present in cell nuclear extracts. The CoRec approach is built upon the PBM technology for the high-throughput measurement of protein-DNA binding^33–35^ and extends work from our lab using PBMs to analyze the DNA binding of TF-COF complexes from cell extracts.^30–32^ In CoRec, we apply nuclear extracts to a DNA microarray and profile the recruitment of a target COF to thousands of customized DNA sequences representing different TF binding sites. We infer the identity of the TFs involved in COF recruitment based on the pattern of DNA sites to which the COF is recruited. Therefore, many TF-COF complexes can be characterized simultaneously, providing a rapid and high-throughput assay to identify TF-COF complexes functioning in a cell.

Briefly, nuclear extracts are applied to a double-stranded DNA microarray, and DNA-bound COFs are labeled using fluorescently labeled antibodies (**Figure 1A**). The CoRec microarray contains consensus binding sites that span the DNA binding site specificity of ∼65% of human TFs, allowing us to profile COF recruitment by the majority of human TFs (**Methods**). Critically, for each of the 346 consensus binding sites we also include all single-nucleotide variant (SV) sites on the microarray, defining 346 consensus+SV probe sets on the array (**Figure 1B).** By profiling the differential recruitment of each COF to the consensus+SV probe sets we can define COF “recruitment motifs” that can be matched against large motif databases (e.g., CIS-BP^36^ and JASPAR^37^) to infer the identity of interacting TFs at the level of TF family^31,32^ (**Figure 1C-D)**. The CoRec approach is inherently multiplexed as COF recruitment to all 346 TF sites is assayed in parallel. Eight individual COFs can be profiled per microarray, allowing thousands of potential TF-COF complexes to be profiled in a single experiment.

**Figure 1:**
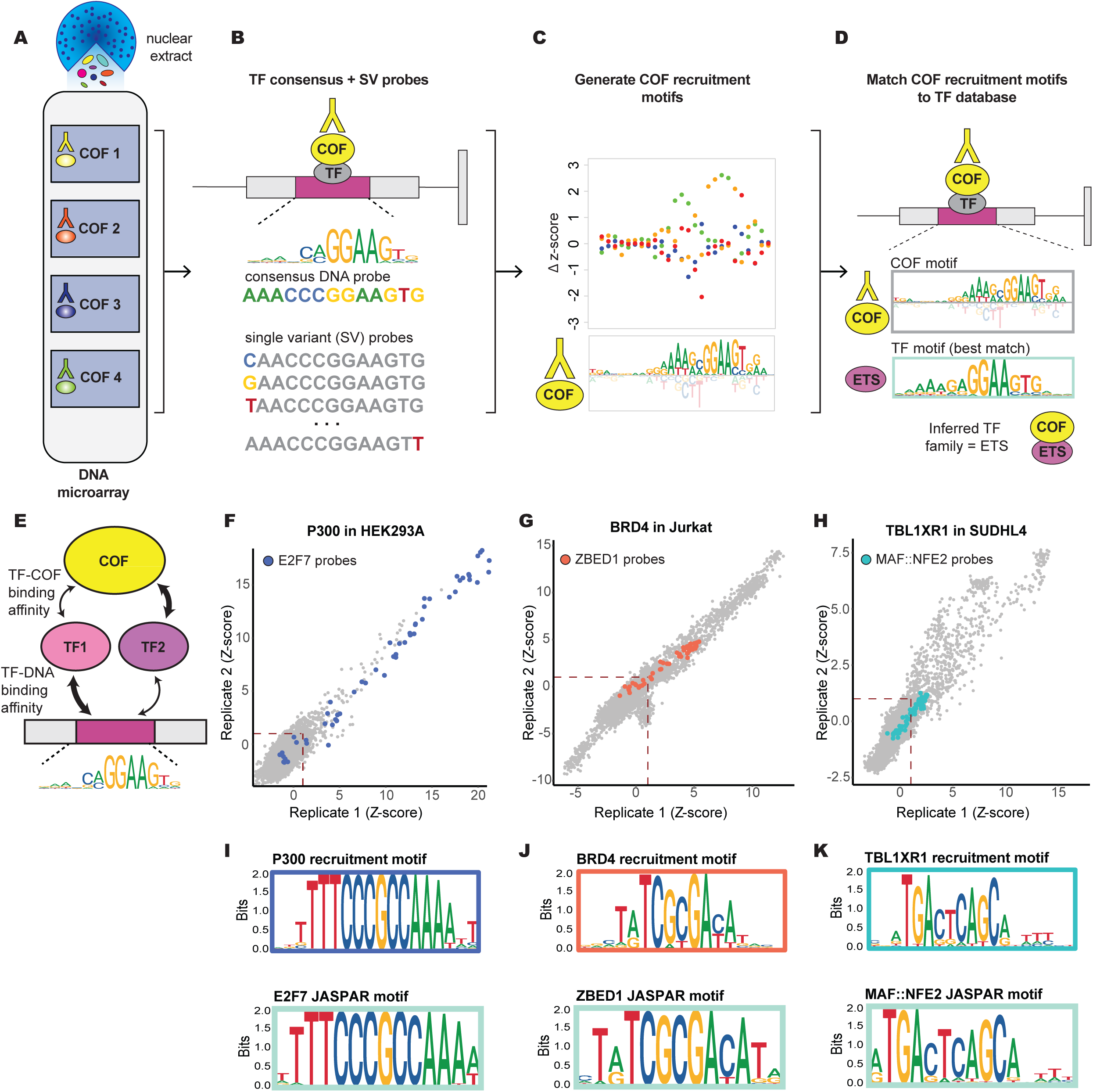
CoRec is an array-based method to profile cell-specific TF-COF interactions. Overview of the CoRec Method. (A) Nuclear extract from cell type of interest is applied to DNA microarray. (B) Microarray contains 346 sets of consensus DNA probes (canonical TF binding site) and all single variant (SV) probes. (C) COF recruitment to all consensus+SV probes sets is used to determine COF recruitment motifs. (D) COF recruitment motifs are matched to TF DNA-binding motifs to infer TF identities. (E) Schematic illustrating that multiple TFs with different COF interaction strengths may contribute to COF recruitment on the microarray. (F-H) Comparison of replicate experiments for three COFs profiled in three cell types. COF recruitment is quantified using microarray fluorescence-based z-scores. Consensus+SV probes associated with select TFs are highlighted. (I-K) COF recruitment motifs corresponding to consensus+SV probes highlighted in each scatter plot are shown along with best-match TF motifs from the JASPAR database.

COF recruitment to each motif is quantified by a *motif strength* calculated from the PBM fluorescence intensities for each consensus+SV probe set. The motif strength depends on TF concentration, TF-DNA binding affinity, and TF-COF interaction affinity (**Figure 1E**). As such, motif strength provides an aggregate measure of COF recruitment by all TFs in the cell that bind to a given DNA motif. For example, the ETS motif strength quantifies the aggregate ability of the ETS family members present in the extract to bind that motif and recruit a given COF. Consequently, the CoRec-defined motif strength represents a cell-specific recruitment activity for individual TF DNA binding-site motifs.

To evaluate the applicability and reproducibility of CoRec, we performed experiments for the COFs P300 (acetyltransferase), BRD4 (scaffold protein) and TBL1XR1 (subunit of NCOR/SMRT repressor complexes) using extracts from three different cell lines - HEK293 (embryonic adrenal precursor cells),^38^ Jurkat (T-cell leukemia cells), and SUDHL4 (B-cell lymphoma cells). Replicate experiments showed excellent agreement in the COF recruitment to all consensus and SV probes (**Figures 1F-H**). The measured COF recruitment strength is quantified by a z-score based on the probe fluorescence values (**Methods**). To demonstrate the sensitivity of CoRec in quantifying differential COF recruitment to DNA variants across a range of binding strengths, we highlight COF recruitment to consensus+SV probe sets derived from motifs for E2F7, ZBED1, and MAF:NFE2 (heterodimer) (**Figure 1F-H**). Despite the wide range of z-scores across the indicated probe sets, the COF recruitment motifs are in excellent agreement with published TF motifs (**Figure 1I-K**) and allow us to infer the identity of TF-COF complexes binding with different strengths across a range of DNA sites (**Figure 1E**). These results demonstrate that CoRec is a robust method to investigate a broad range of TF-COF complexes present in cell extracts.

### CoRec identifies cell-specific TF-COF interaction networks

To evaluate our ability to define cell-specific TF-COF interaction networks, we examined recruitment of the COFs P300, BRD4, and TBL1XR1 to our full CoRec microarray using extracts from three cell types. The TF-COF interaction network is represented as a heatmap indicating significant matches of COF motifs to TF motifs (**Figure 2A**, **Methods**). As many TFs bind DNA with similar sequence specificity, we collapsed TFs into 184 TF specificity ‘clusters’ based on the similarity of their known binding motifs (**File S1, Methods**). Many TF clusters correspond to conventional TF families (e.g. *NFKB_REL* cluster); however, some TF families were split or combined based on motif similarity (e.g. *MYBL_OVOL* is a combined cluster). The intensity of each cell in the heatmap corresponds to the maximum COF motif strength for that TF cluster and represents how strongly a COF is recruited to DNA by TFs in that cluster.

**Figure 2:**
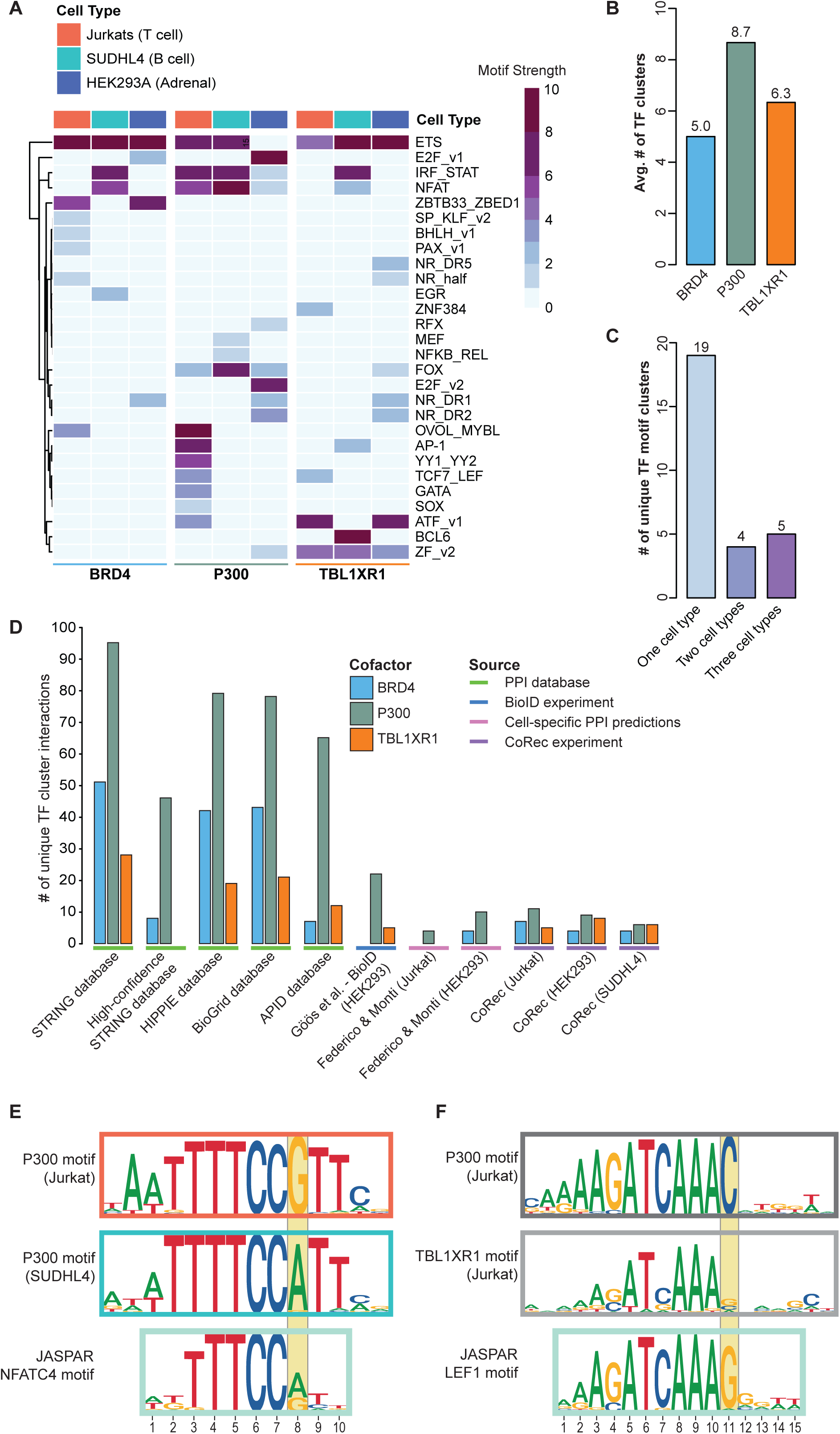
CoRec identifies cell-specific TF-COF interaction networks. (A) Heatmap illustrating CoRec TF-COF interaction data. Columns indicate COFs profiled in each experiment. Rows indicate TF motif clusters recruiting each COF. Heatmap cells are colored by the maximum motif strength of COF motifs matching each TF cluster. (B) Average number of TF clusters identified for COF. (C) Number of unique TF clusters identified in CoRec experiments for different cell types. (D) Number of TF clusters identified for COFs in public PPI databases, individual studies, and CoRec experiments. (E) Comparison of P300 recruitment motifs that matched the NFAT motif cluster found in Jurkat (Top) and SUDHL4 (Middle), and a representative NFAT motif from JASPAR. (F) Comparison of P300 (Top) and TBL1XR1 (Middle) recruitment motifs in Jurkat that matched TCF7_LEF motif cluster, and a representative TCF/LEF family motif from JASPAR.

We identified 28 distinct TF clusters that recruit the three COFs (P300, BRD4, and TBL1XR1) across the three cell types. Comparing the average number of TF clusters per COF, we find the most diverse recruitment for P300 (8.7 clusters), then TBL1XR1 (6.3 clusters), and finally BRD4 (5.0 clusters) (**Figure 2B**). P300 is a broad transcriptional activator with defined sub-domains that interact with a variety of TFs.^39,40^ It has been suggested that P300 must interact with multiple TFs to stabilize its own binding at chromatin, and thus interacting with a large number of TFs may be necessary for P300’s function.^41^ While BRD4 can directly interact with TFs, contributing to site-specific recruitment, it also contains two bromodomains that bind acetylated histones.^42–44^ As our assay involves short (60-bp) un-chromatinized DNA probes, it is of note that BRD4 has the lowest average number of recruitment clusters, consistent with a model in which BRD4 recruitment *in vivo* is mediated by interactions with both TFs and chromatin.

To determine whether the number of CoRec-defined TF-COF interactions is comparable to other approaches, we compared our results to published PPI datasets (**Figure 2D**). To evaluate TF-COF interactions in a single cell type, we first examined data from cell-specific PPI predictions^45^ and proximity-labeling approaches.^46^ We find comparable numbers of TF interactions per COF between CoRec (average of 6.3 clusters per COF) and these published approaches (average of 3 and 9 clusters per COF for cell-specific PPI predictions^45^ and BioID^46^ respectively). To understand how our approach compared to larger datasets, we examined the number of TF-COF interactions reported for each COF in public PPI databases (STRING (v12.0, physical subnetwork)^47^, HIPPIE (v2.3)^48^, BioGrid (release 4.4.221)^49^, APID(version: March 2021)^50^) (**Figure 2D**). These databases aggregate PPI data measured using different approaches (e.g., yeast-two hybrid, co-immunoprecipitation, etc.) and cell types. As expected, the number of PPIs reported for each COF in the databases is higher than the number of CoRec detected interactions given their broad inclusion criteria. However, it was surprising that our CoRec data for TBL1XR1 in only three cell types captured half of the reported interactions in the largest database (14 clusters from CoRec, 28 in STRING), suggesting that interactions in TBL1XR1 are undersampled in existing databases. Despite the large number of PPIs currently represented in the public databases, 26% (12/46) of the COF-TF cluster interactions we identified have not been previously reported (**Figure S1A**). Specifically, we identified five unreported interactions each for TBL1XR1 and BRD4, which have the fewest reported interactions in the databases, and two new TF interactions for P300.

Of the 28 TF clusters that recruit the three COFs, we found that 68% (19/28) were observed in only one cell type, 14% (4/28) were found in two cell types, and 18% (5/28) were observed across all three cell types (**Figure 2C**). The most broadly recruiting TF cluster is the *ETS* cluster, which recruits multiple COFs in all three cell types. ETS TFs comprise a large family with 28 members in humans.^51^ ETS factors regulate house-keeping genes in a number of cell types.^52,53^ Further, they can bind redundantly to regulate house-keeping genes and genes that are constitutively expressed at high levels.^52,54,55^ Examining the STRING PPI database, we find that the *ETS* cluster has the most interactions (718 individual ETS-COF interactions) of any of our reported TF clusters, supporting their broad COF interaction ability. Our data suggests that promiscuous COF recruitment across cell types is a feature of the *ETS* cluster and that more generally this may be a feature of TFs that regulate constitutively expressed genes.

In contrast to the broadly-acting TF clusters, cell type-specific interactions can highlight cell-specific TF functions. For example, the *MEF* and *BCL6* clusters are cell-type specific and only recruit COFs in SUDHL4 B cells. BCL6 and MEF2B (member of the *MEF* cluster) are both crucial regulators of B cell development, and changes in their COF interactions have been implicated in diffuse large B-cell lymphoma (DLBCL).^56–59^ In our dataset, BCL6 interacts strongly with TBL1XR1. Notably, the BCL6-TBL1XR1 interaction is rarely disrupted in germinal center B cell (GCB)-DLBCL, the subtype to which SUDHL4 belongs, but is often disrupted in the more aggressive DLBCL subtype known as activated B cell (ABC)-DLBCL.^59^ These results support the ability of CoRec to identify functionally relevant TF-COF interactions in a cell type-specific manner. We further note that because CoRec assays endogenous proteins, it provides a straightforward method to profile the impact of cell-specific mutations on TF-COF interactions and DNA binding, as exemplified by the BCL6-TBL1XR1 interactions in DLBCL subtypes.

### TF-COF binding motifs can differ from canonical TF binding motifs

A central feature of CoRec is that recruitment motifs are determined independently for each COF. Comparison of COF motifs can reveal both COF and cell-specific differences that suggest additional means for achieving gene regulatory specificity. For example, the *NFAT* cluster recruits P300 in both Jurkat and SUDHL4 cells, but the recruitment motifs differ between the cell types. In Jurkat cells, the P300 recruitment motif reveals a strong preference for a guanine at position 8 (5’-NNTTTCC**<u>G</u>**NN-3)’, whereas in SUDHL4 cells there is a strong preference for adenine at this position (5-NNTTTCC**<u>A</u>**NN-3)’ **(Figure 2E**). NFAT family member motifs appear to have a relatively equal preference for A and G at this position and previous studies have noted NFATC2 binding to both of these types of sequences, albeit with varying affinities.^60^ Strikingly, in SUDHL4 cells BRD4 and TBL1XR1 are also recruited to NFAT motifs with the same preference for adenine at position 8, suggesting that is a cell-type specific difference (**Figure S1B**).

COF-specific differences within individual cell types were also observed. For example, TBL1XR1 and P300 are recruited to the *TCF7_LEF* cluster in Jurkat cells, but the motifs for each COF differ in distinct ways. For P300, position 11 of the motif shows a distinct preference for cytosine, whereas for TBL1XR1 there is little or no base preference at this position (**Figure 2F**). In contrast, the *TCF_LEF* cluster member motifs all indicate a strong-to-moderate preference for guanine at this position (represented by the LEF1 motif for illustration). While the weak preference shown for TBL1XR1 may result from differences in motif generation, the strong cytosine preference for P300 suggests an altered specificity for TF-P300 complexes. Cell-specific COF motifs identified by CoRec provide a means to identify additional mechanisms of TF-COF binding specificity.

### Gene expression, protein levels, and PPI data do not predict TF-COF interactions

To investigate whether mRNA levels, protein levels, and PPI data can explain our observed cell-specific TF-COF interaction data, we further analyzed our CoRec results from Jurkat and SUDHL4 cell lines for which we have both RNA-seq and whole proteome mass-spectrometry data.^61,62^ The COFs themselves have similar mRNA and protein levels in both cell types (**Figure S1C-D**), suggesting that cell-specific TF-COF differences are not a result of COF levels. Focusing on the TFs, we identified 16 TF clusters that recruit COFs in only one of the two cell types (**Figure 3A**). We then evaluated whether at least one TF associated with each of the 16 TF clusters was differentially expressed (log2(FC) >2) on the mRNA or protein level, which could explain the observed cell-specific difference in COF recruitment (**Figure 3A, Tables S2-S3**). We found that in 75% (12/16) of the cases, there was at least one TF cluster member that had differential mRNA that might explain the observed COF recruitment differences. Examining proteomic data, we find that 63.5% (10/16) of the differential COF recruitment could be explained by changing TF protein levels. Finally, we asked whether PPI data from the STRING database would further support predictions of differential recruitment. Of the cases where altered TF-COF interactions were supported by differential mRNA or protein levels, we found 50% (8/16) were also supported by PPI data for the relevant COF, highlighting the difficulty in predicting how TF-COF networks may change across cell types.

**Figure 3:**
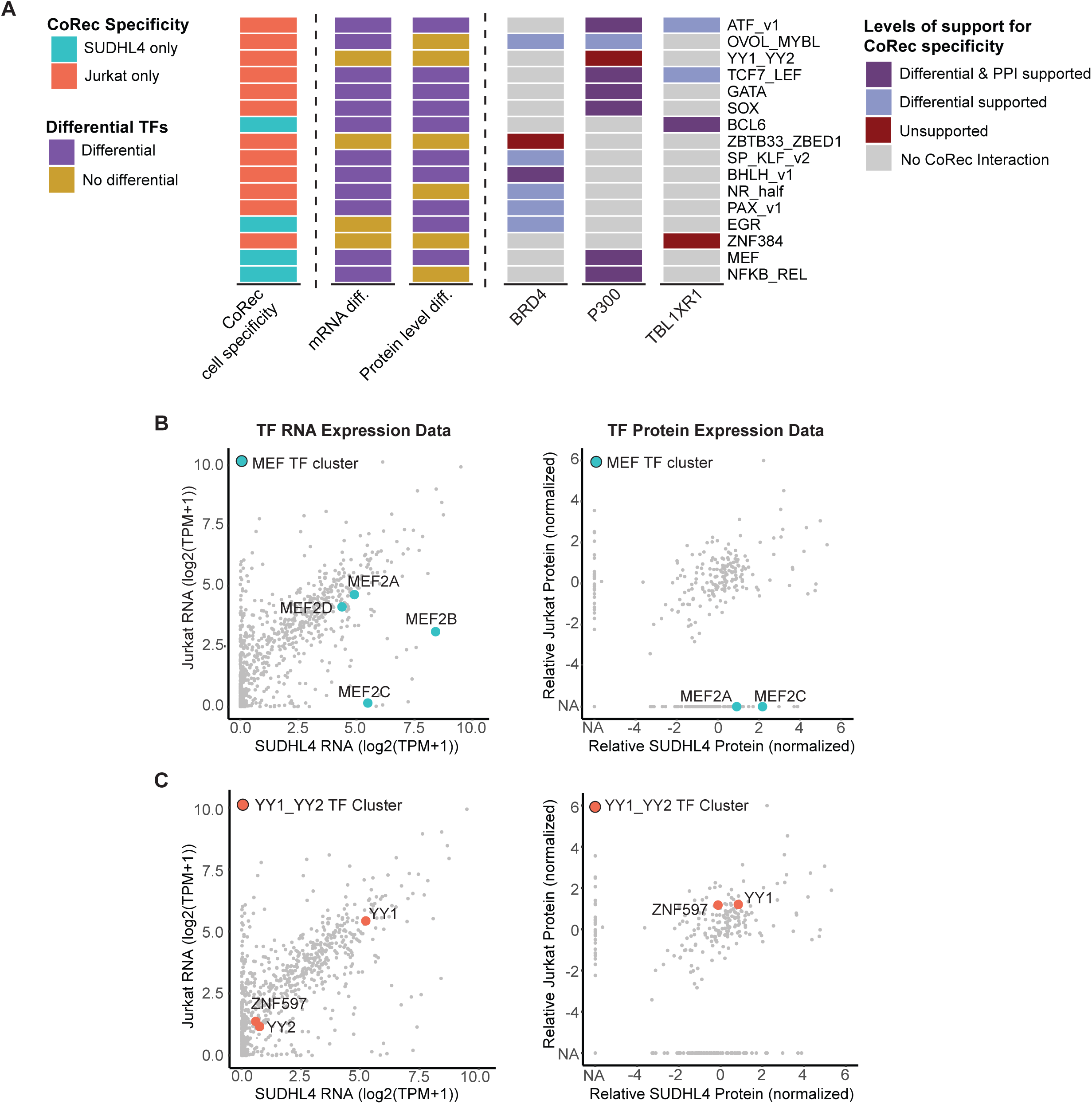
Gene expression, protein levels, and PPI data cannot predict cell-specific TF-COF interactions. (A) Comparison of CoRec cell specificity to cell-specific mRNA expression, protein levels, and PPI data. (Column 1) Cell-specific recruitment observed for different TF clusters. (Columns 2,3) Differential expression or protein levels (log2(FC) > 2) for at least one member within each TF cluster that is in the same sense as the cell-specific COF recruitment. (Columns 4,5,6) Levels of support for COF cell-specificity. ‘Differential’ indicates support from either differential expression or protein levels. ‘PPI’ indicates the indicated COF can interact with at least one of the differential TF for that cluster. (B-C) Comparison of mRNA expression (left) and protein levels (right) for TFs in Jurkat and SUDHL4 with members of *MEF* cluster or *YY1_YY2* cluster highlighted.

Two examples illustrate how we can leverage expression, proteomic, and PPI datasets to refine our CoRec measurements and also highlight the importance of cell-specific measurements. The *MEF* cluster recruits P300 in SUDHL4 but not in Jurkat (**Figure 2A**). Members of the *MEF* cluster are expressed in both cell types, but MEF2B and MEF2C are expressed at higher levels in SUDHL4 (**Figure 3B**). Examining the protein levels themselves the trend is even more striking as no *MEF* members are identified in Jurkat while MEF2A and MEF2C proteins are both detected in SUDHL4. Furthermore, interactions between P300 and both MEF2A and MEF2C have been reported^47^; therefore, a plausible scenario is that differences in MEF2 TF levels could explain the observed recruitment differences we see between SUDHL4 and Jurkat. In contrast, *YY1_YY2* recruits P300 in Jurkat but not in SUDHL4; however, TFs in this cluster show no differences in mRNA or protein levels (**Figure 3C**), suggesting that P300 recruitment is likely regulated by post-translational modifications or other mechanisms that differ between these two cell types. These examples demonstrate the difficulty in explaining (or predicting) differential recruitment data using mRNA, proteomic, and existing PPI data, and highlight the need for cell-specific measurements, as afforded by CoRec.

### CoRec identifies cell state-dependent TF-COF interactions

Cell-specific profiling of TF-COF complexes provides a means to examine the TF-COF complexes driving epigenetic changes as cells differentiate and respond to signals. Histone tail acetylation has been correlated with cis-regulatory element (CRE) activity and gene expression changes.^63–65^ Therefore, we sought to determine whether profiling KAT-TF interactions could identify transcriptional activators important for a given biological output. We profiled seven COFs from three main KAT subfamilies (P300/CBP family, GCN5/PCAF family, and MYST family) to examine recruitment of KATs in resting and T cell-receptor (TCR)-stimulated Jurkat T cells. As transcriptional changes downstream of TCR signaling have been found to occur in as little as one hour,^66^ we examined the early changes in KAT recruitment (45 min post stimulation) that likely initiate the gene expression response.

We observed 111 TF-COF interactions across the KATs and T cell conditions, involving 43 unique TF motif clusters (**Figure 4A**). Of the 111 interactions, 20% (22/111) of the interactions are gained upon T cell stimulation, 15% (17/111) are lost upon stimulation, and 65% are present in both conditions, demonstrating that even at this early 45-min time point, ∼35% of the TF-COF interactions have been altered. We identified several broad categories based on KAT recruitment, including 4 TF clusters with strongly induced recruitment, 2 TF clusters with partially induced recruitment, 3 TF clusters with diminished recruitment, and 15 TF clusters with recruitment specific to one KAT. All KATs, except GCN5, exhibited changes in TF interactions in response to T cell signaling at this time point (**Figure 4B**). Of the KATs that showed signal induced TF cluster interactions, TIP60 exhibited the largest number of recruiting clusters (8 clusters), while MOF had the fewest (1 cluster). Comparing the number of KAT-interacting TF clusters to those from cell-specific BioID measurements (HEK293 cells), we observed comparable numbers of interactions, i.e., an average of 15.9 clusters per COF for CoRec versus 11.8 per COF for BioID (**Figure S2A**). The data highlights that even at relatively short stimulation times, we observe large-scale changes in the KAT-TF network of T cells.

**Figure 4:**
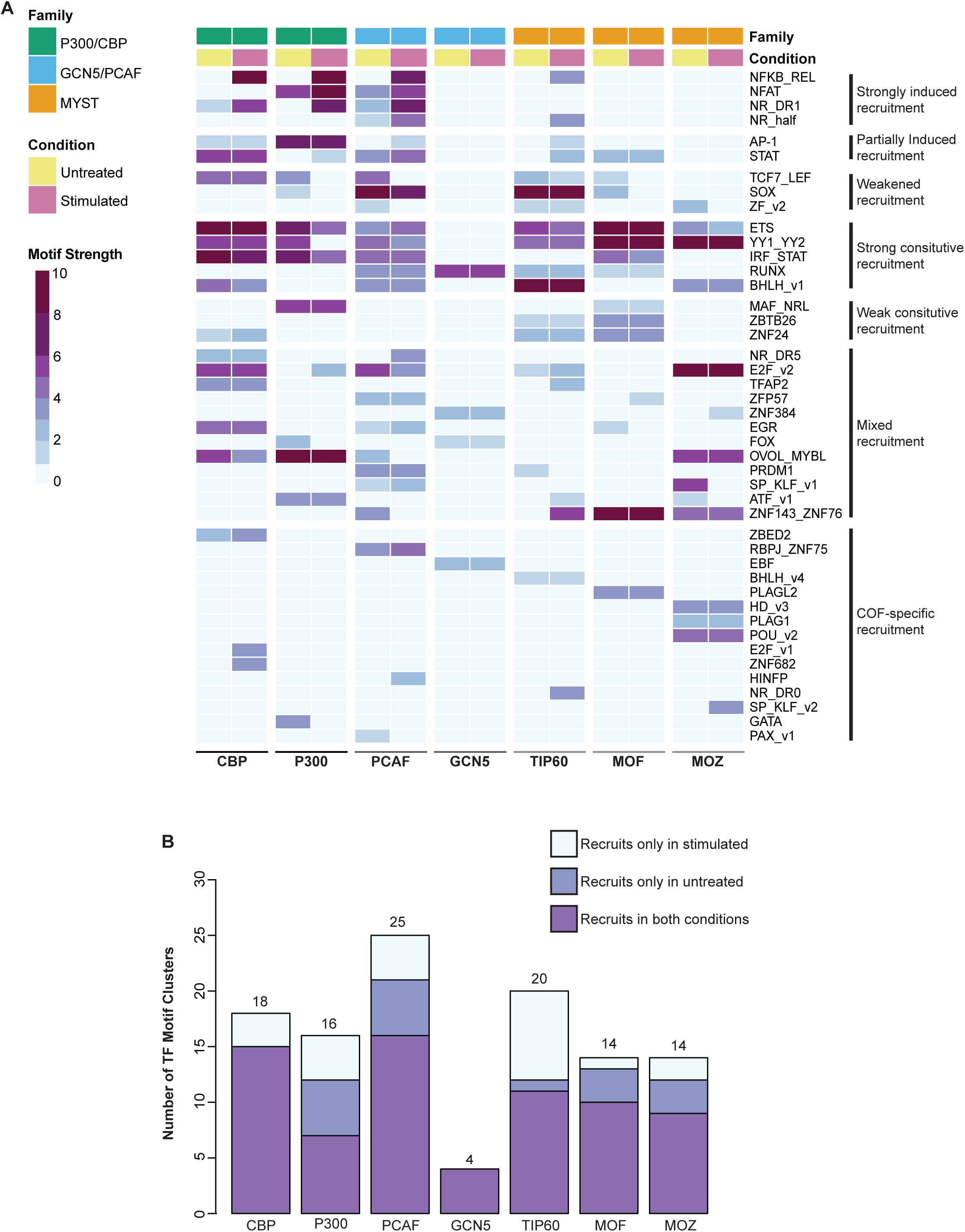
KAT-TF interaction network reveals KAT and TF specificities. (A) Heatmap of KAT-TF interaction network. Columns indicate profiled KATs and experimental conditions: untreated Jurkat T cells or 45 min post TCR stimulation. Rows indicate TF motif clusters recruiting each COF. Heatmap cells are colored by the maximum motif strength of COF motifs matching each TF cluster. (B) Total number of TF clusters recruiting each KAT and condition specificity.

### Stimulus-dependent activators recruit multiple KATs

Canonical downstream activators of TCR and CD28 co-receptor signaling involve three major TF pathways: NF-κB, NFAT, and AP-1.^67,68^ Interactions between KATs and members of these TF families have been reported to contribute to gene activation.^47,69–74^ To determine whether profiling KAT-TF interactions can identify these TF activators, we identified TF clusters that recruited multiple KATs more strongly after T cell stimulation (i.e., motif strengths are strictly higher after activation, Δz-score > 1.5) (**Figure 4A**). Four TF clusters met this criteria – *NFKB_REL*, *NFAT*, *NR_half*, *NR_DR1* – two of which match the expected canonical TCR pathways (NF-κB and NFAT). The motifs identified for *NFKB_REL* and *NFAT* are consistent with motifs of RelA and NFATC1, regulators known to be activated by TCR/CD28 signaling (**Figure 4A**). Relaxing our activator selection criteria to allow for both constitutive and induced KAT recruitment we identified two additional clusters, *AP-1* and *STAT*. The *AP-1* cluster represents the third canonical TCR pathway, and recruits PCAF and TIP60 upon stimulation, but constitutively recruits CBP and P300 (**Figure 4A**). This differential activator status for AP-1 suggests a more complex KAT interaction landscape than the strict stimulus-induced activators NF-κB and NFAT.

Among our set of strict activators we also identified two Nuclear Receptor TF clusters. The *NR_half* cluster contains motifs defined by a single 5’-AGGTCA-3’-type NR half-site motif, while the *NR_DR1* cluster contains motifs defined by direct repeats of the half-site separated by a one nucleotide spacer (DR1). The NR4A family binds the *NR_half* motifs, and all members of this family have been shown to be rapidly induced upon TCR stimulation including as early as 30 min post-stimulation.^75–77^ NR4A1 has also been shown to interact with PCAF,^78^ and we find an increased recruitment of PCAF to the *NR_half* cluster upon stimulation (**Figure 4A**). The *NR_DR1* cluster represents motifs bound by a range of type II NRs.^79,80^ However, peroxisome proliferator-activated receptors (PPARs) TF family bind as heterodimers with Retinoid X receptor α (RXRα) nuclear receptor to *NR_DR1* cluster motifs,^79^ and both the PPAR family and RXRα play a role in T cell activation, regulating cytokine expression and contributing to T cell survival.^81,82^ We note that the *NR_half* and *NR_DR1* clusters recruit distinct repertoires of KATs, supporting the conclusion that separate NR complexes bind to each motif. These results demonstrate that profiling KAT interactions can provide a protein-level approach to identify transcriptional activators of cell signaling events.

### Promiscuous KAT recruitment is a feature of transcriptional activators in T cells

KATs catalyze deposition of different repertoires of histone acetylation marks^64,83^ (**Figure 5A**), many of which can facilitate, or are correlated with, enhanced transcription.^84,85^ Genome-wide maps of histone 3 lysine 27 acetylation (H3K27ac) and P300 binding are widely used to predict enhancer activity^86–88^; therefore, we anticipated that TF activators may interact strongly with P300 (and paralog CBP) following TCR stimulation. However, the simple combination of H3K27ac marks and P300 binding does not identify all enhancers,^89–91^ and recruitment of other KATs (e.g. PCAF, GCN5, TIP60, MOZ, MOF) has been observed at active regulatory elements.^63,84,89,92^ These observations support a model in which many KATs are recruited to active regulatory elements^84^; however, it is not clear if this is by individual TFs each recruiting separate KATs, by promiscuous KAT recruitment by TFs, or by a combination of recruitment by TFs and other chromatin-associated factors (e.g., chromatin marks or other regulatory COFs).

**Figure 5:**
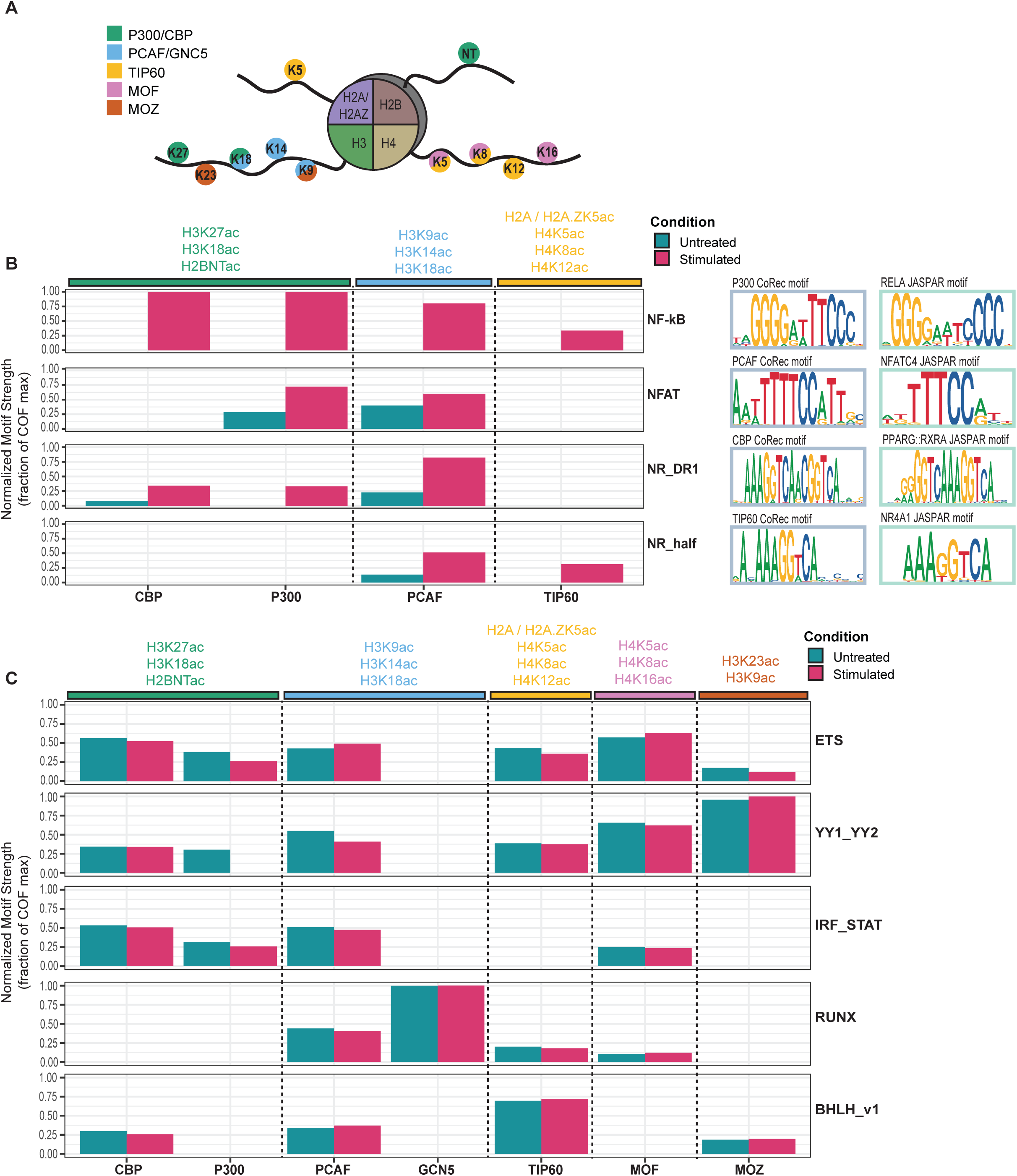
KAT-TF interactions reveal promiscuous KAT recruitment. (A) Schematic indicating histone lysine residues acetylated by different KATs. (B) COF recruitment of four TF clusters with TCR-stimulation induced recruitment. COF recruitment is quantified by a normalized motif strength relative to the max for each KAT. Histone acetylation marks catalyzed by each KATs are indicated. Representative COF motifs matching each TF cluster are shown along with matching TF motifs from JASPAR. (C) COF recruitment of constitutively recruiting TF clusters. COF recruitment is quantified as in (B).

In light of these findings, we re-examined the KAT recruitment specificity of both stimulus-dependent and constitutive activators. Focusing first on strict stimulus-induced TF activators (*NFKB_REL*, *NFAT*, *NR_DR1*, *NR_half*), we find induced recruitment of paralogs CBP and P300 that deposit H3K27ac, as well as induced recruitment of PCAF and TIP60 that deposit different histone marks (**Figure 5B**). When also considering AP-1, the other canonical TCR signaling TF pathway, we find that stimulus-enhanced PCAF recruitment was the one common feature across all stimulus-induced TF activators. PCAF is known to deposit H3K9ac and H3K14ac, both marks that have been correlated with active regulatory elements.^91^ Notably, we did not find a wide-spread stimulus-dependent increase in recruitment for MOZ, MOF, GCN5 (**Figures 4A-B**). These observations demonstrate that stimulus-specific TF activators downstream of TCR signaling interact with multiple KATs in a signal-dependent fashion that would predict deposition of a number of histone acetylation marks at activated regulatory elements.

Studies have identified binding of multiple KATs at active transcriptional regulatory elements in resting T cells.^63,84^. Therefore, we next examined constitutive recruiters in our dataset to assess the repertoires of KATs recruited. We identified five TF clusters that exhibit constitutive, promiscuous recruitment (>4 KATs with motif strength >3 in both conditions): *ETS, YY1, IRF_STAT, RUNX,* and *bHLH* (**Figure 5C**). TFs from these families are known to regulate either house-keeping genes (ETS, YY1),^52,93,94^ or genes associated with T cell functions (ETS, RUNX, bHLH, IRF_STAT).^95–99^ For example, IRF1 and IRF4 act as competing pioneer factors in T cells,^95^ and members of ETS and RUNX were found to occupy the vast majority of accessible chromatin regions in mouse CD4 and CD8 T cells.^96^ These data demonstrate that broadly-acting T cell activators are associated with, and can be identified by, constitutive KAT recruitment. Furthermore, the observation that KATs are promiscuously recruited to constitutive activators provides a mechanism for the wide-spread co-occupancy observed for KATs and multiple histone acetylation marks at regulatory elements in T cells.^63,100^

### CBP and P300 paralogs exhibit unique KAT recruitment profiles

CBP and P300 are paralogous KATs with homologous TF interacting domains.^101^ Despite their sequence homology and co-occupancy at many genomic loci,^63^ studies have identified functional differences between them.^102–104^ Strikingly, while we find a number of TFs that recruit both CBP and P300, we find numerous differences in their recruitment profiles. There are 10 TF clusters that recruit both P300 and CBP, 8 that uniquely recruit CBP and 6 that are unique to P300 (**Figure 4A**). However, even for the 10 shared TF clusters, 6 exhibit different patterns of condition-specific recruitment. For example, the *E2F_v2* cluster recruits CBP in both resting and stimulated Jurkat cells but recruits P300 only in the stimulated condition. TFs in this cluster are implicated in controlling the proliferation of T cells after TCR stimulation and have been shown to recruit both P300 and CBP to activate target genes.^105^ The interaction data indicate joint recruitment of CBP and P300 upon stimulation, but a CBP-specific role in resting T cells. Overall, our data demonstrate that CBP and P300 exhibit differential recruitment profiles in both resting and TCR-stimulated T cells, providing a possible mechanism for their overlapping yet specific functions.

### KAT-TF interactions highlight ascertainment bias in PPI databases

To evaluate potentially novel KAT-TF interactions in our dataset, we compared our results to PPI data in the STRING database (**Methods**).^47^ This database contains both direct and indirect associations between proteins, and includes PPIs determined from experiments as well as computational predictions such as text mining or conserved co-expression.^47^ Evaluating KAT-TF interactions at the level of TF clusters, we find that 40% (44/111) of our KAT-TF interactions represent previously unreported interactions (**Figure 6A**). We identified at least one unreported TF cluster interaction for every KAT except for P300. Strikingly, the majority of interactions with MOF and MOZ were unreported (85.7% and 92.9%, respectively). This was in contrast to P300 and CBP, where most TF cluster interactions had been previously reported (**Figure 6B**). To evaluate whether the number of novel interactions we find might be explained by the extent to which a KAT has been previously studied, we compared the number of PPIs for each KAT in the STRING database to the percent of known KAT-TF interactions in our data (**Figure 6C**). We identified a clear relationship (logarithmic fit, adjusted R2 = 0.92, p-value <0.01) that suggests that the number of unreported interactions identified for each KAT is likely a result of ascertainment bias of PPIs reported in the public databases. These data indicate that interactions with certain KATs are understudied, and suggest that similar biases may be true for other COFs.

**Figure 6:**
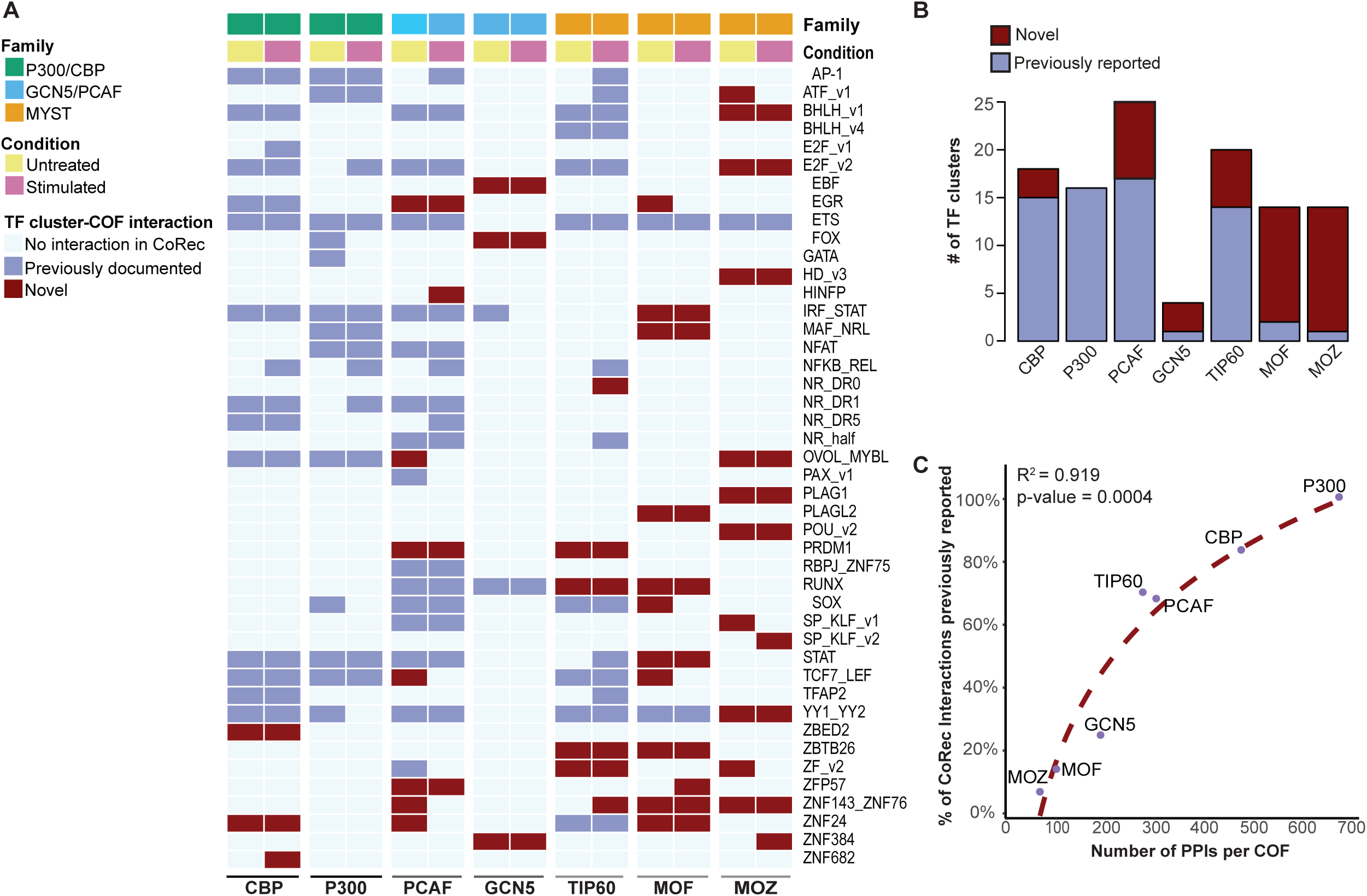
CoRec provides a means to profile TF recruitment of understudied KATs. (A) Heatmap indicating novel KAT-TF interactions. Columns indicate profiled KATs and experimental conditions: untreated Jurkat T cells or 45 min post TCR stimulation. Rows indicate TF motif clusters recruiting each COF. Heatmap cells are colored to indicate whether PPIs with each KAT and a member of the TF clusters have been reported in a public PPI database. (B) Proportion of KAT-TF cluster interactions that are novel. Interactions annotated as in (A). (C) Percentage of KAT-TF interactions reported in PPI databases compared to the total number of PPIs reported for that KAT.

### KAT recruitment by NF-κB correlates with H3K27ac levels at induced genes

We found that TF activators downstream of TCR signaling gained interactions with KATs by 45 min post stimulation. To determine whether changes in histone acetylation are observed at this early time point, and whether they relate to changes in KAT-TF interactions, we performed ChIP-seq for H3K27ac marks in resting and TCR-stimulated (45 min) Jurkat T cells. We identified a total of 25,481 H3K27ac ChIP-seq peaks across both conditions; with 150 peaks gaining H3K27ac post-stimulation (FDR < 0.01, log2(FC) > 2), and 27 peaks losing acetylation (FDR < 0.01, log2(FC) < -2) **(Figure 7A**). A representative locus of the known TCR-induced gene *RelB,* showed induced acetylation peaks (**Figure 7D**); Gene Ontology (GO) analysis of genes associated with induced peaks returns terms related to stimulated T cells including “T cell activation” (**Figures S3A-S3B**). To relate changes in acetylation with gene expression, we compared genes that were associated with increased acetylation (log2(FC) > 2) to genes that did not change (log2(FC) < 0.5 & > -0.5) or were reduced (log2(FC) < -2). Genes with induced acetylation were expressed at lower levels prior to stimulation (**Figure 7B**) and at higher levels post stimulation (**Figure 7C**); these genes included many known activated T cell genes, including *NR4A1*, *EGR3*, and *IL2RA* (encoding CD25) (**Figure 7A**).

**Figure 7:**
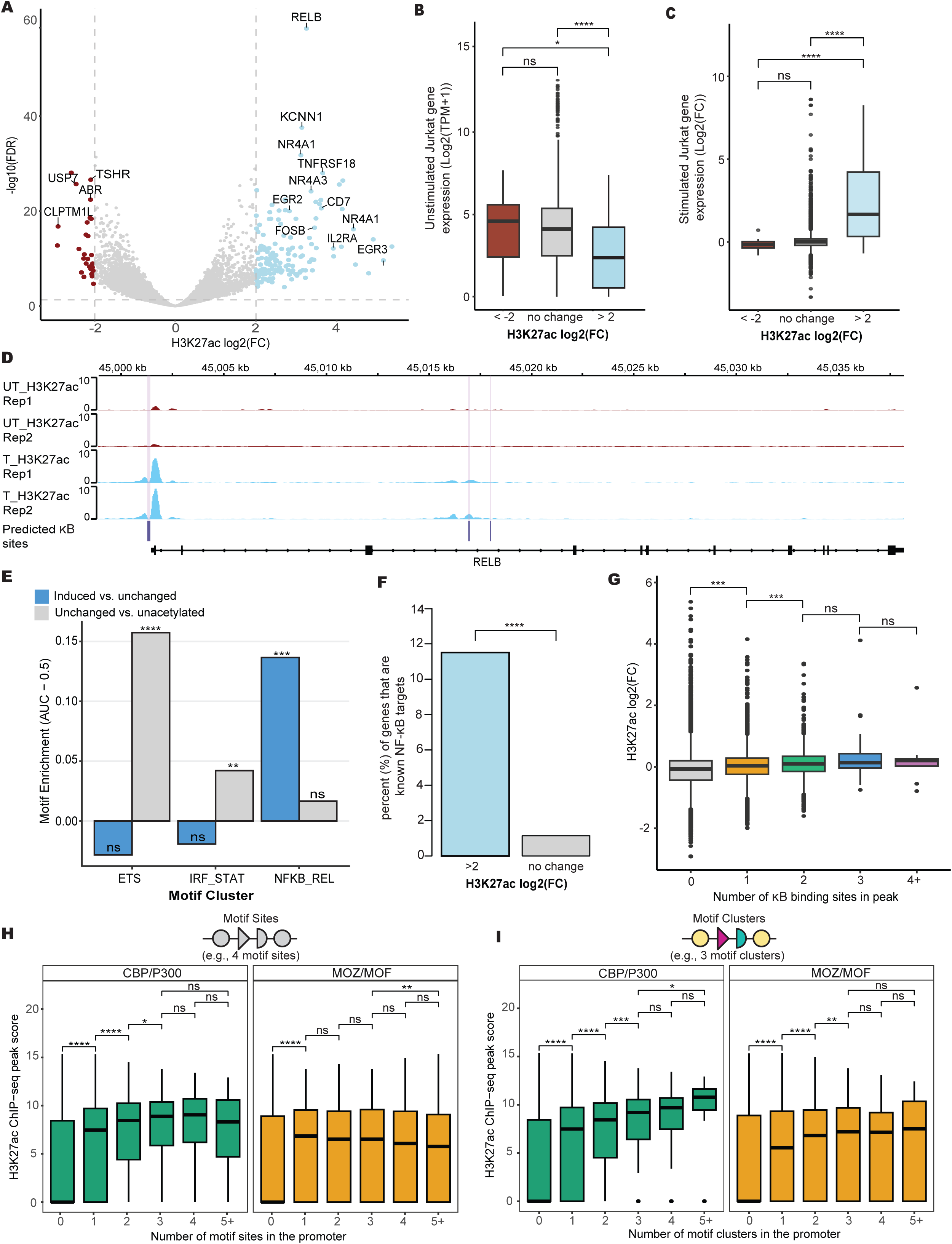
P300/CBP recruitment identifies TFs associated with genome-wide H3K27ac levels. (A) Differential H3K27ac peaks in TCR-stimulated Jurkat T cells. Log2(fold change) of H3K27ac levels between untreated and 45 min post TCR-stimulation conditions. Select genes linked to differential acetylation peaks are indicated. (B) Gene expression levels in unstimulated Jurkat for genes associated with TCR-stimulated decreasing (log2(FC) < -2), unchanging (-0.5 < log2(FC) < 0.5), or increasing (2 < log2(FC)) H3K27ac levels at associated peaks. (C) TCR-stimulated gene expression changes (2 hours post-stimulation) for gene sets defined as in (B) (D) ChIP-seq H3K27ac tracks in untreated (UT_H3K27ac) and 45 min TCR-treated (T_H3K27ac) Jurkat cells. κB binding sites predicted within the region are indicated. (E) Enrichment of motifs from TF clusters in loci defined by differential H3K27ac levels. (F) Proportion of known NF-κB target genes associated with induced H3K27ac. (G) Impact of κB site number on changes in H3K27ac levels at ChIP-seq peaks. (H) Impact of the number of specific COF-recruiting motifs on promoter H3K27ac levels (peak scores). Motifs evaluated in gene promoters (-500 nt to +100 nt from TSS). Data shown for motifs recruiting CBP/P300 (green) and motifs recruiting MOZ/MOF and not CBP/P300 (orange). (I) Impact of the number of COF-recruiting motifs from unique TF clusters (i.e., heterotypic motifs) on promoter H3K27ac levels (peak scores). Motif identity and promoter elements defined as in (H). Adjusted p-values are as follows; ns > 0.05, * is <= 0.05, ** is <= 0.01, *** is <= 0.001, **** is <= 0.0001.

As H3K27ac marks are deposited by P300 and CBP, we examined whether motifs from TF clusters with enhanced CBP/P300 recruitment following stimulation were enriched in the promoters that gained acetylation. We found that NF-κB motifs, those associated with the strongest induced recruitment of both CBP and P300, were the only significantly enriched motifs in these promoters compared to promoters that did not gain acetylation (**Figure 7E**). In contrast, motifs from the two strongest constitutive recruiters of P300 and CBP, the *IRF_STAT* and *ETS* clusters, were not enriched in induced genes but were enriched in promoters with constitutive acetylation. Further supporting the role of NF-κB in altering acetylation levels and promoting gene activation, genes associated with induced H3K27ac peaks are enriched in NF-κB target genes(**Figure 7F**).^106^ For example, within the cis-regulatory regions of of the NF-κB target gene *RelB* there are several regions of increased acetylation that overlap κB binding sites, and previous studies have indicated that the *RelB* transcription is dependent on NF-κB RelA binding (**Figure 7D**).^107^ To further evaluate the impact of NF-κB sites on H3K27ac marks, we examined whether the number of predicted κB binding sites that overlapped with the H3K27ac peaks correlated with H3K27ac levels. We found a statistically significant increase in H3K27ac levels with a single NF-κB site, and a further increase with two sites, but no further gain beyond two sites (**Figure 7G**). These results suggest that the stimulus-enhanced CBP/P300 interactions seen for NF-κB factors correspond to concomitant gains in histone acetylation and induced genes expression.

### Heterotypic clusters of KAT-recruiting TF sites correlate with promoter H3K27ac levels

We identified a number of TF clusters that constitutively recruit KATs (**Figure 4A**) and hypothesized that they likely play a role in histone acetylation levels in both resting and stimulated T cells. To test this hypothesis, we examined whether the presence of TF motifs associated with constitutive CBP/P300 recruitment correlated with constitutive genome-wide H3K27ac levels at gene promoters. We first examined whether multiple instances of the CBP/P300-recruiting motifs predict higher promoter acetylation levels, and we found significantly higher levels of promoter acetylation with the occurrence of up to three motifs (**Figure 7H**). As a control, we examined the impact of motifs that do not recruit CBP/P300 but do recruit MOZ/MOF, and we found no increase beyond one motif for this group. Given that CBP and P300 were recruited by several TF clusters, we then asked whether heterotypic motif groups - defined by motif matches from *different* TF clusters - were also predictive of acetylation levels. Strikingly, we found an even stronger trend in which promoters with motifs from up to 5 or more CBP/P300-recruiting TF clusters predicted increasingly higher acetylation levels. This trend was not seen with motifs from TF clusters that recruit only MOZ/MOF and not CBP/P300 (**Figure 7I**). This trend of acetylation levels with heterotypic motif clusters was also stronger than seen for homotypic motif clusters (**Figure S3C**). These results demonstrate that KAT-TF networks can determine TF families that are predictive of genome-wide acetylation levels, and that high levels of promoter acetylation is likely a result of heterotypic clusters that contain binding sites for multiple KAT-recruiting TFs from distinct families.

## Discussion

CoRec is a method for rapid profiling of cell-specific TF-COF complexes, and provides an alternative to other methods for high-throughput analysis of PPIs that focuses on DNA-bound TF-COF complexes. In CoRec, TFs that recruit target COFs are inferred using DNA binding motifs. As multiple TFs can often bind the same motif, our assay provides an aggregate, quantitative measure of COF recruitment by TFs that bind the same DNA sites (**Figure 1E**). We demonstrate that CoRec can be used to identify cell-specific TF-COF interactions for different classes of COFs (**Figure 2A**). Profiling KAT-TF interactions in resting and stimulated Jurkat T cells, we found that 35% of the interactions were condition specific, highlighting the importance of cell context for TF-COF interactions. We show that profiling interactions with KATs provides a means to identify transcriptional activators, and propose that using CoRec to profile interactions with other classes of COFs can provide a means to characterize TF function. CoRec can be readily applied to different cell systems, requiring only nuclear extracts and cofactor antibodies for labeling. We anticipate that using CoRec in different cell states and disease contexts (e.g., different cancer cell types) will provide insights into how TF-COF networks alter chromatin state and gene regulation.

Investigating regulators of histone acetylation, we find that promiscuous recruitment of KATs is a prominent feature in our KAT-TF network in T cells, with 66% (29/44) of TF clusters recruiting at least two KATs and 30% (13/44) recruiting at least four KATs (**Figure 4A**). The recruitment of multiple different KATs to the same DNA sites, whether by single TFs or by paralogous TFs that bind the same DNA sites, predicts co-occurrence of KATs and their cognate histone acetylation marks at regulatory elements. Indeed, studies in primary T cells that examined genome-wide binding of five KATs (P300, CBP, PCAF, MOF, TIP60) ^63^ and the occurrence of 18 acetylation marks ^100^ reported wide-spread colocalization of these KATs and histone acetylation marks. This wide-spread co-occurrence suggests low KAT specificity for many genes, which is at odds with genetic perturbation data for individual KATs.^84^ These observations led Anamika *et al.* ^84^ to propose a model in which initiation of gene expression is mediated by specific KATs, whereas maintenance of expression requires less KAT specificity and involves the recruitment of many KATs to the acetylated chromatin environment. As KAT complexes often contain bromodomains, the authors proposed that KAT co-occurrence may result from bromodomain-mediated interactions with acetylated histone tails.^84^ While our data is consistent with this model, it suggests that the KAT co-occurrence at regulatory elements is also the result of promiscuous KAT recruitment by constitutive TF activators. Beyond the mechanisms by which multiple marks are deposited at regulatory elements, it also remains unclear what roles these multiple acetylation marks play for maintenance of gene expression. One potential explanation is that dynamic TF-COF interactions ^108^, in combination with active deposition and removal of acetylation marks ^109^, make multiple acetylation marks necessary in order to maintain open chromatin and create a more constant level of expression. Future studies that integrate TF-COF interaction data with gene perturbation approaches should help to clarify the roles of constitutive TF activators on promiscuous KAT recruitment and the impact of different KATs on the maintenance of basal levels of gene expression.

The combinatorial action of multiple TFs is a feature of eukaryotic enhancers and promoters; however, deciphering the individual roles of TFs within regulatory elements remains a challenge.^110^ Here, we sought to understand the role of TFs by examining their recruitment of COFs associated with specific epigenetic marks. We focused on H3K27ac marks associated with the activity of promoters and enhancers. To characterize potential activators in T cells, we profiled TFs that recruit the KATs CBP and P300 that deposit H3K27ac. In TCR-activated T cells, we found that the NF-κB family had the strongest stimulus-induced recruitment for both CBP and P300, and that NF-κB family motifs were the most enriched in loci that gained H3K27ac (**Figure 5B and 7E**). These results are consistent with the known role of NF-κB as a potent activator downstream of T cell signaling.^67,68,111,112^ In resting T cells, we identified a number of TF clusters that recruit either CBP or P300 (**Figure 4A**). Unexpectedly, we found that heterotypic clusters of these TF motifs at individual promoters was correlated strongly with the H3K27ac levels (**Figure 7I**), and was more predictive than the total number of TF motifs (**Figure 7H**) or the number of homotypic motifs (**Figure S3C**). We hypothesize that heterotypic motif clusters may be predictive of H3K27ac as both CBP and P300 contain multiple TF-interacting subdomains^101^ that could facilitate synergistic CBP/P300 recruitment.^13,110^ Indeed, a recent study has highlighted the individual roles of these subdomains in P300 genome-wide binding.^41^ It is unclear if similar relationships exist for other histone marks; however, similar studies performed for other classes of COFs should provide insight into how heterotypic motif clusters relate to additional features of promoters or enhancers.

## Acknowledgments

This work was supported by National Institutes of Health (NIH) R01 A116829, NIH R21 HG011289, and NIH RO1 AI51051 to T.S., and a seed grant to T.S. and T.D.G. from Boston University Genome Sciences Institute.

## Author Contributions

M.M.I., J.L.K., T.S. and T.D.G jointly conceived of the study. R.M. and D.B. developed the software to design, analyze, and visualize the CoRec experiments. M.M.I. and H.H. performed the experimental work; J.L.K, R.Z, performed preliminary experiments. M.M.I., R.M., H.H., and T.S., analyzed and interpreted the experimental results. M.M.I., R.M., and T.S. wrote the manuscript.

## Declaration of Interests

The authors declare no competing interests.

## STAR Methods

### Cell culture and nuclear extractions

SUDHL4 (CRL-2957), Jurkat E6-1 (TIB-152), and HEK293 (CRL-1573) cells were obtained from ATCC. SUDHL4 and Jurkat cells were grown in suspension RPMI 1640 Glutamax media (Thermofisher Scientific, Catalog #72400120) with 10% heat-inactivated fetal bovine serum (Thermofisher Scientific, Catalog #132903) and Jurkat cells medium was also supplemented with 1mM sodium pyruvate (Thermofisher Scientific, Catalog #16140071). Cells were grown at 37 °C with 5% CO2. T175 (CELLTREAT, Catalog #) non-treated flasks were used when culturing SUDHL4 and Jurkats cells for experiments. HEK293 cells were grown in low glucose DMEM (Cytiva, Catalog # SH30021) supplemented with 10% FBS at 37 °C with 5% CO2. HEK293 cells were grown in cell culture treated T225 flasks (CELLTREAT, catalog #) for experiments.

Jurkat cells intended to be stimulated as well as paired untreated cells were plated at 1 x 10^6^ cell/ml 2 hours prior to stimulation. To stimulate selected cells, soluble anti-CD28, CD3 & CD2 (ImmunoCult™ Human CD3/CD28/CD2 T Cell Activator, Catalog #10970) was added to media at manufacturers recommended 25 ul/ml. Cells were agitated slightly and placed back in the incubator for 45 minutes. After 45 minutes, both the untreated and stimulated cells were processed either for nuclear extraction or ChIP-seq.

### Nuclear extraction

The nuclear extract protocols are as previously described ^30,32^ with modifications detailed below. Approximately 100 million cells were harvested for each nuclear extraction protocol. To harvest suspension cells, the cells were collected in a falcon tube and placed on ice. Suspension cells were pelleted by centrifugation at 500xg for 5 min at 4ºC. Supernatant was collected and cell pellets were resuspended in 10 mL of 1X PBS with protease inhibitor and pelleted again at 500xg for 5 min at 4ºC and PBS wash was aspirated leaving behind the cell pellet. To collect HEK293 cells for nuclear extraction, media was aspirated from the flask and 10 ml of warmed 1x PBS was used to gently wash the cells. PBS was aspirated off and 10 ml of ice-cold 1x PBS with protease Inhibitor was added to the flasks. A cell scraper was used to lift the HEK293 cells from the flask and cells were collected in a falcon tube and put on ice. Collected cells were pelleted at 500xg for 5 min at 4ºC before PBS was aspirated. All remaining steps were performed the same between suspension and HEK293 cells. To lyse the plasma membrane, the cells were resuspended in 1.5 mL of Buffer A (10mM HEPES, pH 7.9, 1.5mM MgCl, 10mM KCl, 0.1mM Protease Inhibitor, Phosphatase Inhibitor (Santa-Cruz Biotechnology, Catalog #sc-45044), 0.5mM DTT (Sigma-Aldrich, Catalog #4315) and incubated for 10 min on ice. After the 10 min incubation, Igepal detergent (final concentration of 0.1%) was added to the cell and Buffer A mixture and vortexed for 10 s. To separate the cytosolic fraction from the nuclei, the sample was centrifuged at 500xg for 5 min at 4ºC to pellet the nuclei. The cytosolic fraction was collected into a separate microcentrifuge tube. The pelleted nuclei were then resuspended in 100 µL Buffer C (20mM HEPES, pH 7.9, 25% glycerol, 1.5mM MgCl, 0.2mM EDTA, 0.1mM Protease Inhibitor, Phosphatase Inhibitor, 0.5mM DTT, and 420mM NaCl) and then vortexed for 30 s. To extract the nuclear proteins (i.e., the nuclear extract), the nuclei were incubated in Buffer C for 1 h while mixing at 4ºC. To separate the nuclear extract from the nuclear debris, the mixture was centrifuged at 21,000xg for 20 min at 4ºC. The nuclear extract was collected in a separate microcentrifuge tube and flash frozen using liquid nitrogen. Nuclear extracts were stored at -80ºC.

### CoRec experiments

Microarray DNA double stranding and PBM protocols are as previously described. ^30,32,33,113^ Any changes to the previously published protocols are detailed. Double-stranded microarrays were pre-wetted in HBS (20mM HEPES, 150mM NaCl) containing 0.01% Triton X-100 for 5 min and then de-wetted in an HBS bath. Next the array was incubated with a mixture of the binding reaction buffer (20mM HEPES, pH 7.9, 100mM NaCl, 1mM DTT, 0.2mg/mL BSA, 0.02% Triton X-100, 0.4mg/mL salmon testes DNA (Sigma-Aldrich, Catalog #D7656) and nuclear extract for 1 h in the dark. The array was then rinsed in an HBS bath containing 0.05% Tween-20 and subsequently de-wetted in an HBS bath. After the protein incubation, the array was incubated for 20 min in the dark with 20mg/mL primary antibody for the COF of interest (**Table S1**) diluted in 2% milk in HBS. After the primary antibody incubation, the array was rinsed in an HBS bath containing 0.1% Tween-20 and de-wetted in an HBS bath. Microarrays were then incubated for 20 min with 20mg/mL of either alexa488 or alexa647 conjugated secondary antibody (**Table S1**) diluted in 2% milk in HBS. The array was rinsed in an HBS bath containing 0.05% Tween-20 and then placed in a Coplin Jar containing 0.05% Tween-20 in HBS. The array was agitated in solution in Coplin Jar at 125 rpm on an orbital shaker for 3 min and then placed in a new Coplin Jar with 0.05% Tween-20 in HBS to repeat the washing step. It was then placed in a Coplin Jar containing HBS and washed for 2 min as described above. After the washes, the array was de-wetted in an HBS bath. Microarrays were scanned with a GenePix 4400A scanner and fluorescence was quantified using GenePix Pro 7.2. Exported fluorescence data were normalized with MicroArray LINEar Regression.^113^ Analysis of normalized CoRec microarray data was performed using the publicly available CoRec analysis package (https://github.com/Siggers-Lab/CoRec).

### CoRec microarray design

Non-redundant TF binding position-weight matrix (PWM) models from the JASPAR 2018 core vertebrate set were obtained using the JASPAR2018 R bioconductor package. The 452 human models were collapsed into consensus sequences and filtered for equivalence based on nucleotide identity and relative sequence length (cutoff > 0.9). Sequence length filter was used to ensure that composite site models for two TFs (e.g., A + B) and half-site models (e.g., A or B) were both included in the final design. Filtering led to a final set of 346 TF models to be included in the final design. These 346 models represents ∼65% of human TFs based on comparison against human motifs in the CIS-BP database^36^ (motif comparison p-value < 0.0005 using Tomtom (MEME Suite version 5.3.3).^114–116^ TF consensus binding sites were embedded within a 34-nt DNA probe sequence attached to a 24-nt common primer sequence. For all 346 consensus sequences DNA probes corresponding to each possible single-nucleotide variant (SV) sequence were also included on the microarray (**Figure 1B**). The 60-nt probe sequences were organized as follows: 2-nt GC cap + 34-nt binding site (TF site or SV probe) + 24-nt common primer. 261 background target DNA probes were included in the design to estimate background fluorescence intensities in the experiments. Background probes sequences were randomly selected from the human genome (hg38).

### CoRec motif generation and motif strength

Log-transformed PBM fluorescence values (median fluorescence over 5 replicate probes) were normalized against background fluorescence levels to yield a z-score that quantifies binding to each probe sequence:

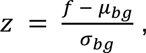

where *f* is the log fluorescence value of the probe, *μ*_*bg*_ is the mean background log fluorescence value, and *σ*_*bg*_ is the standard deviation of the background log fluorescence values. COF binding specificity to each consensus+SV probe set was modeled using a Δz-score binding motif that captures the change in binding for every SV substitution across the consensus binding site. Δz-score motifs are defined as:

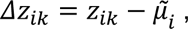

where *z*_*ik*_ is the z-score for nucleotide variant *k* at position *i* of the motif and 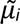 is the median z-score for all nucleotide variants at position *i*. The Δz-score binding motif is akin to a binding energy matrix that quantifies the impact of nucleotide variants to the total binding energy. COF binding strength is quantified using a *motif strength* score based on the median z-score of top-scoring 15% of probes contributing to the motif (i.e., for this consensus+SV probe set) (**Files S4-S5**).

To allow for motif comparison to published PWM-type TF binding models, the Δz-score motifs were also transformed into position probability matrices (PPM) using a Boltzmann distribution formalism and the calculated motif strength (MS) (**Files S6-S7**).

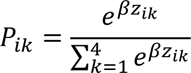

*β* is calculated as follows for each individual CoRec motif:

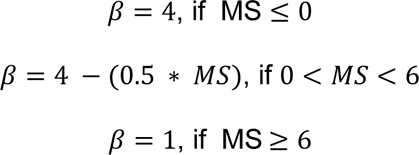

To determine if COF recruitment specificity was conserved between conditions or cell types, we performed pairwise comparison of replicate PPM motifs between each condition. Specifically, for a specific consensus+SV probe set we determined the two replicate CoRec motifs in each condition. We then performed all four pairwise motif comparisons between motifs from the two conditions. If the minimum motif Euclidean distance (i,e., highest motif similarity) was less than 0.25 then we annotated these motifs as conserved across conditions for this consensus+SV probe set (**Files S8-S9**).

### TF motif cluster generation

Motif clusters were generated using a set of 946 human TF binding models from the JASPAR2022^37^ CORE database. Pairwise motif comparison was performed using Tomtom (MEME Suite version 5.3.3^114–116^) with default settings. The distance between two motifs was defined as max(15 +log10(p-value)), 0) using the Tomtom returned p-values. Initial motif clusters were created using agglomerative clustering with the complete linkage method (i.e., clusters were combined based on the maximal distance between two motifs from the different clusters). Clusters were then manually curated to account for the canonical TF families represented (**File S1**). For integration with publicly available PPI datasets, we mapped individual human TFs to our TF motifs clusters. To do this we mapped 6445 human TF motifs in the CIS-BP database to our TF motif clusters. CIS-BP motifs were assigned to their best match (p-value < 0.0005, TomTom default settings). At this stringency, 65% of ∼1200 human TFs can be assigned to TF motif clusters (**File S1**).

### Matching COF motifs to TF binding motifs

To infer the identity of TFs recruiting each COF, CoRec motifs were matched to databases of TF binding motifs. First, we filtered out low-quality CoRec motifs based on a minimum motif strength of 0.4 and a minimum average information content of 1.0 in any window of 5 consecutive positions. Non-reproducible motifs were also removed. Reproducible motifs were defined as motifs that passed the motif strength and information content thresholds in at least two independent replicates and that had an inter-replicate PPM motif Euclidean distance less than 0.4. CoRec motifs meeting these thresholds were then compared to the set of 946 motifs used to generate the motif clusters using the memes R package (version 1.2.5),^117^ an R wrapper of the MEME Suite (version 5.3.3).^114^ The comparisons were done with the run_tomtom() function using Euclidean distance and requiring a minimum overlap of 5 nucleotides between the target and query motifs (dist = “ed”, min_overlap = 5). A match was considered significant if at least one replicate CoRec motif had an adjusted p-value less than 0.01. Adjusted p-values were calculated by Tomtom^115,116^ as the raw p-value multiplied by the number of motifs in the reference library. Each CoRec motif was assigned to the motif cluster corresponding to its best matching reference motif. If replicate motifs were assigned to different clusters, the motif(s) with poorer p-values were reassigned to the cluster of the COF recruitment motif with the best match p-value.

### RNA-seq, proteomic and PPI analysis

<u>RNA-seq analyses:</u> To examine gene expression values in resting Jurkat and SUDHL4 cells, RNA-seq data was obtained from the DepMap consortium.^62^ To examine induced gene expression in TCR/CD28-activated Jurkat T cells we used data from Felce et al.^118^ Differential expression analysis was performed using EdgeR package (Release (3.18)).^119^ Preprocessed expected count profiles for each cell type were used with as input, with filters for 100 reads required for each gene, and a bcv value set to 0.2. Genes with log2(FC) greater than 2 or less than -2 were determined to be differentially expressed (**File S2**). <u>Proteomic analyses:</u> To examine protein levels across cell types, normalized relative protein expression levels for resting Jurkat and SUDHL4 were obtained from the DepMap consortium and used as provided.^81^ Proteins with log2(FC) greater than 2 or less than -2 were determined to be differentially expressed between the cell lines (**File S3**). <u>PPI analyses:</u> To compare our CoRec-defined interactions with PPI data from public databases, individual TF-COF interactions in PPI databases were mapped onto TF motif clusters annotations using 6445 human TF motifs in CIS-BP (described above). Using this mapping we could compare COF interactions with TF motif clusters across PPI and CoRec datasets.

### ChIP-seq experiments

Jurkat cells in complete growth medium at concentration of 1 x 10^6^ cells/ml were collected in a conical tube. For each sample, 4×10^7^ cells were crosslinked with 1% formaldehyde (final concentration) (Thermo Fisher, cat #033314.AP) for 10 min at RT with gentle rotation. Crosslinking was stopped by adding 125 mM final concentration of glycine solution in PBS and cells were rotated at RT for 5 min. Fixed cells were pelleted at 1000xg for 10 min at 4°C and washed twice with 14 ml of ice cold PBS and pelleted at 1000xg for 10 min at 4°C each time. Washed cell pellet was resuspended in 1 ml of ice cold PBS and transferred to 1.5 ml lo-bind DNA tube (Eppendorf, Catalog # 022431005) and centrifuged at 2400xg for 5 min at 4°C and flash frozen with liquid nitrogen and stored at -80°C for later use.

Soluble Chromatin was prepared as previously described^120^ with some modifications (outlined below). Frozen cell pellets were thawed and resuspended in 1.2 ml of Lysis Buffer 1 (50 mM Hepes, 140 mM NaCl, 1 mM EDTA, 10% Glycerol, 0.5% Igepal, 0.25% Triton X-100, 0.1mM Protease Inhibitor), nutated for 15 min at 4°C, and pelleted at 10000xg for 5 min at 4°C. The same procedure was repeated with lysis buffer 2 (200 mM NaCl, 1 mM EDTA, 0.5 mM EGTA, 10 mM Tris-HCL, 0.1mM Protease Inhibitor) at room temperature followed by pelleting at 10000xg for 5 min at 4°C. Pellets were washed with 400 ul of sonication buffer (0.1% SDS in TE) and then resuspended in 1.4 mL of sonication buffer. Resuspensions were then split into two 1.5 ml lo bind DNA tubes for sonication (each tube with 700 ul of liquid). During sonication each tube was placed in a 1.5 ml microfuge tube placed in Benchtop 1.5 ml Tube Cooler (Active Motif, cat # 53076). The nuclei were sonicated using Active Motif Q120AM sonicator with a 2 mm Probe (Active motif cat # 53056) at 40% amplitude for 40 min with 30 s ON and 30 s OFF cycles (80 cycles total). Cell debris was pelleted at 21000xg for 10 min at 4°C. Soluble chromatin was transferred to a new 1.5 ml lo bind DNA tube, 300 ul of sonication buffer was added along with additional reagent to create RIPA buffer (150 mM NaCl, 0.1% NaDOC, 1% Triton X-100, 5% glycerol). 30 μl of the combined soluble chromatin was saved to be checked for chromatin shearing upon reverse-crosslinking via 1.5% agarose gel and Bioanalyzer 2100 using the DNA High Sensitivity Kit (Agilent, Catalog #5067-4626). The rest of the soluble chromatin mixture was mixed end-over-end agitation for 10 minutes at 4°C then flash frozen and stored at -80°C.

For immunoprecipitation, each chromatin sample was thawed on ice 2 hours prior to experiment and precleared with 20 ul of washed Protein A/G dynabeads via incubation for 1 hour on a rocking platform at 4°C. Precleared chromatin was separated from beads via spin at 12000xg for 10 min at 4°C and supernatant was transferred to new 1.5 ml lo bind DNA tubes. DNA concentration was then measured via Qubit dsDNA HS Assay Kit (Invitrogen, Catalog # Q32851). For immunoprecipitation, 25 ug of precleared chromatin and 3ug rabbit polyclonal anti-H3K27ac antibody (Abcam, ab4729) was mixed together and nutated overnight at 4°C. The next day 14.5 ul of washed protein A dynabeads beads was added to each sample and the mixture was nutated for 4 hours at 4°C. After 4 hours, samples were placed on the magnet and supernatant was gently removed. 700 ul of wash buffer 1 (150 mM NaCl, RIPA buffer) was added to the sample and nutated for 15 minutes before samples were placed on magnet and supernatant was removed. This process was repeated for wash buffer 2 (400 mM NaCl, RIPA buffer), wash buffer 3 (TE, 250 mM LiCl, 0.5% NaDOC, 0.5% NP40), and wash buffer 4 (TE, 0.02% Triton). After removing wash buffer 4, beads were resuspended in elution buffer (TE, 250 mM NaCl, 0.3% SDS) supplemented with 0.8 U of proteinase K and moved to 0.2 mL PCR tubes. The samples were then uncrosslinked in the thermocycler for 42°C for 30 min, 65°C for 5 hours, 15°C for 10 min. ChIP DNA was then purified using the Qiagen MinElute Reaction Cleanup Kit (Catalog #28204) and eluted in 12 ul of 1X TE buffer. Samples concentrations were measured via Qubit and library prep was performed using the NEBNEXT Ultra II DNA library prep kit for Illumina (NEB, catalog #E76455S) following the provider’s instruction manual. Amplified libraries were Bioanalyzed again to check the library preparation success. Samples were pooled by molarity and sequencing on NovaSeq 6000-S4 to obtain ∼35 to 40 million reads per sample.

### ChIP-seq analysis

Raw sequencing reads were trimmed with Trimmomatic v0.36 (SLIDINGWINDOW:4:20 MINLEN:20).^121^ Trimmed reads were then aligned to the human genome (GRCh38, Ensembl release 102^122^) using Bowtie2 v2.4.2 (--very-sensitive-local -X 2000).^123^ H3K27ac peaks were identified separately for resting and TCR-stimulated Jurkat cells using Genrich v0.6 (-r -y -a 100 -E human_excluded_regions.bed,human_scaffolds_chrM.bed) (https://github.com/jsh58/Genrich). The human_excluded_regions.bed file was downloaded from the ENCODE project (accession: ENCFF356LFX). The human_scaffolds_chrM.bed file contains the coordinates of the human genome scaffolds and the full mitochondrial chromosome. Reproducible peaks were identified using Genrich’s automatic collective peak calling approach to combine replicates. Replicates were also analyzed separately for downstream DiffBind analysis.

To determine differential peaks H3K27ac between the untreated and the TCR-stimulated samples we use the R package DiffBind (Release 3.4.11).^124^ Replicate bam read and peak files were used as inputs. Samples were normalized and differential peaks were called using defaults with comparison being called between the treatment groups. DiffBind calculated log(FC) and FDR values were used to determine differential peaks between the two sets of samples. All DiffBind H3K27ac peaks were linked to genes using R package ChIPseeker (v1.30.3)^125^ using GRCh38, Ensembl release 102 ^122^ and org.Hs.eg.db (v3.8.2)^126^ with tssRegion determined as - 3000 to 3000 (default setting). If a gene had multiple peaks in which some changed and some did not, it was called an induced or reduced gene and was filtered out of the “no change” group.

### Motif analysis

Motifs were identified in genomic regions (e.g., H3K27ac peaks or gene promoters) using FIMO (--max-strand) (version 5.3.3).^114,115^ Regions were scored using -log10(p-value) of the single best occurrence of each motif. If a sequence had no occurrences of a given motif with a p-value <= 1e-5, the sequence was assigned a score of 0 for that motif. To assign scores to TF motif clusters (as opposed to single motifs), all motifs associated with the TF cluster were evaluated and the maximum motif score was used. Motif enrichment between sets of genomic elements (e.g., promoters with induced acetylation relative to unchanged promoters) was quantified using the two-sided Wilcoxon rank-sum test.

For motif analyses of gene promoters, we defined promoters as the region from 500 bp upstream to 100 bp downstream of the TSS for a protein coding gene (GRCh38, Ensembl release 102). Alternative TSSs were not considered. Acetylated promoters were defined as induced if they overlapped DiffBind regions (log2(FC) >= 1.5), or as unchanging if they overlapped DiffBind regions (log2(FC) <= 0.1). Promoter acetylation level was defined as the maximum Genrich score (i.e., -log10(p-value)) for any overlapping H3K27ac ChIP-seq peak from either condition. Unacetylated promoters were identified as promoters that overlapped an ATAC-seq peak but not an H3K27ac peak. ATAC-seq peaks were defined using publicly available data in resting Jurkat cells (GEO accession: GSM4706085) and identified using Genrich v0.6 with ATAC-seq mode enabled (-j). Promoters that were unacetylated in both conditions were assigned an acetylation score of 0.

## Supplemental Titles and Legends

**Supplemental Figure 1: Profiling TF-COF interactions provides information on cell-specific TF-COF interactions and TF-COF complex binding motifs.**

(A) Heatmap indicating CoRec TF-COF interactions that have been previously reported in the STRING database.

(B) COF recruitment motifs in SUDHL4 that match the NFAT cluster and a representative consensus NFAT motif from JASPAR. SUDHL4-specific nucleotide preference highlighted in yellow.

(C-D) mRNA expression or normalized protein levels for all COFs for unstimulated Jurkat and SUDHL4 with specific COFs profiled highlighted.

**Supplemental Figure 2: Profiling KATs using CoRec produced the expected number of TF-COF interactions.**

(A) Number of TF clusters identified for KATs profiled in public PPI databases, individual studies, and CoRec experiments.

**Supplemental Figure 3: Integration of CoRec with other datasets can provide additional insights.**

(A-B) Top 10 GO enrichment categories for genes associated with stimulus-induced increase in H3K27ac levels.

(C) Impact of the max number of COF-specific motifs (i.e., homotypic motifs) on promoter H3K27ac levels (peak scores). Motifs evaluated in gene promoters (-500 nt to +100 nt from TSS). Data shown for motifs recruiting CBP/P300 (green) and motifs recruiting MOZ/MOF and not CBP/P300 (orange).

(D) Impact of the number of COF-recruiting motifs from unique TF clusters (i.e., heterotypic motifs) on gene expression levels. Motif identity and promoter elements defined as in (C).

**Table S1: Antibodies used for CoRec experiments.**

**File S1: TF motifs and assigned clusters.**

**File S2: Differential mRNA expression data for unstimulated Jurkat and SUDHL4 cells.**

**File S3: Differential protein level data for unstimulated Jurkat and SUDHL4 cells.**

**Files S4-S5: CoRec-generated z-score motifs.**

**Files S6-S7: CoRec-generated PPMs.**

**Files S8-S9: CoRec-generated motifs.**

## Notes

### Competing Interest Statement

The authors have declared no competing interest.

