## Supplementary figures and images for "Rapid profiling of transcription factor-cofactor interaction networks reveals principles of epigenetic regulation"

### Supplemental Figures 1-3

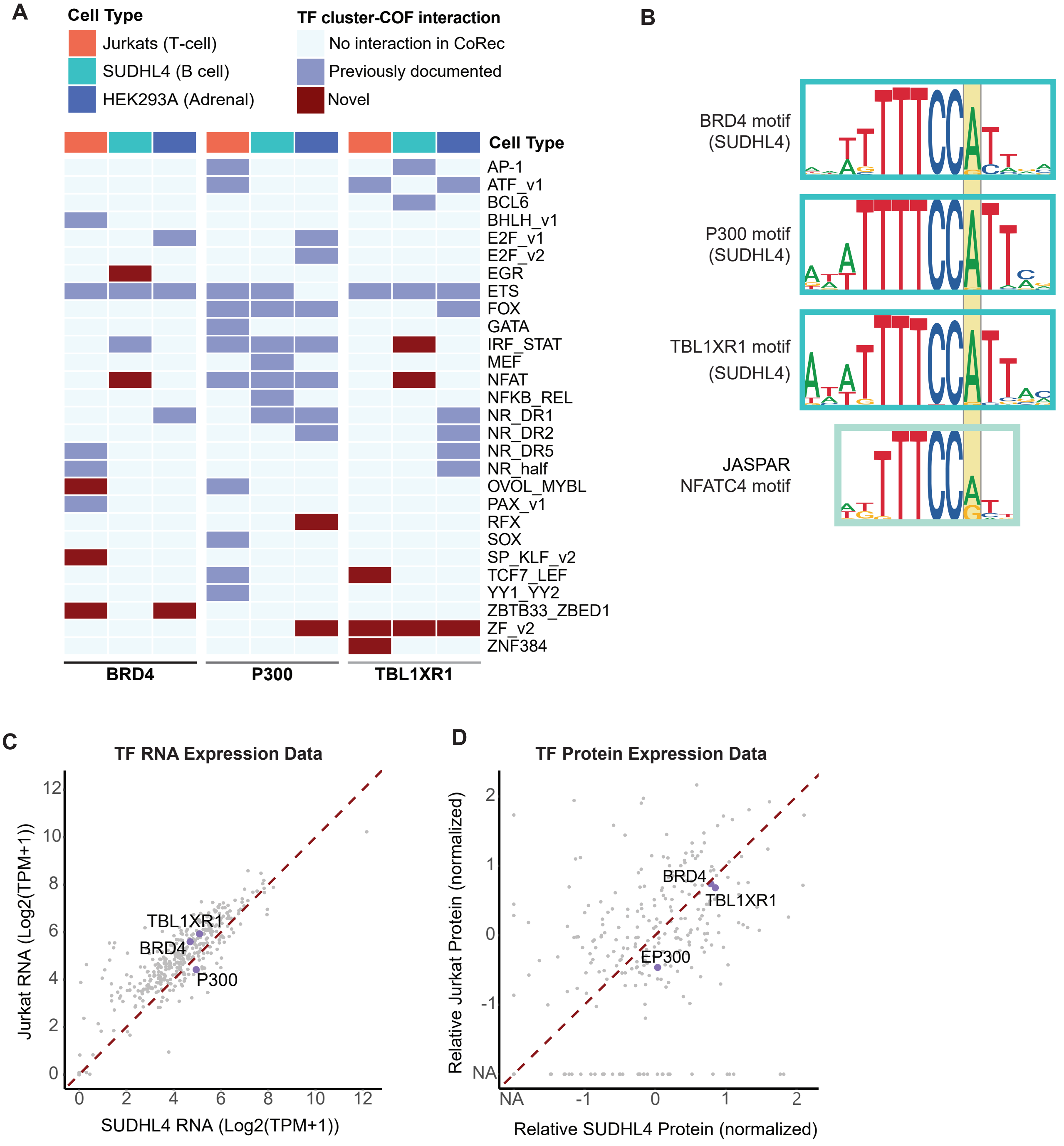

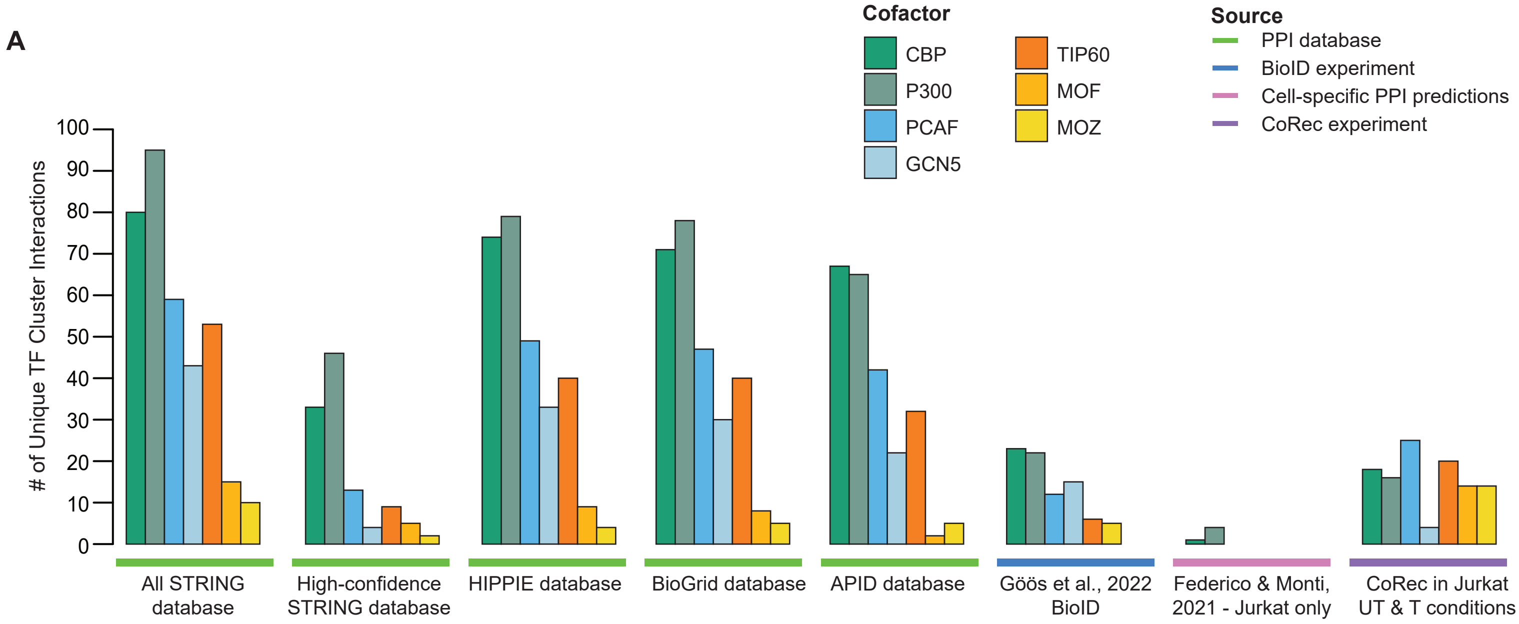

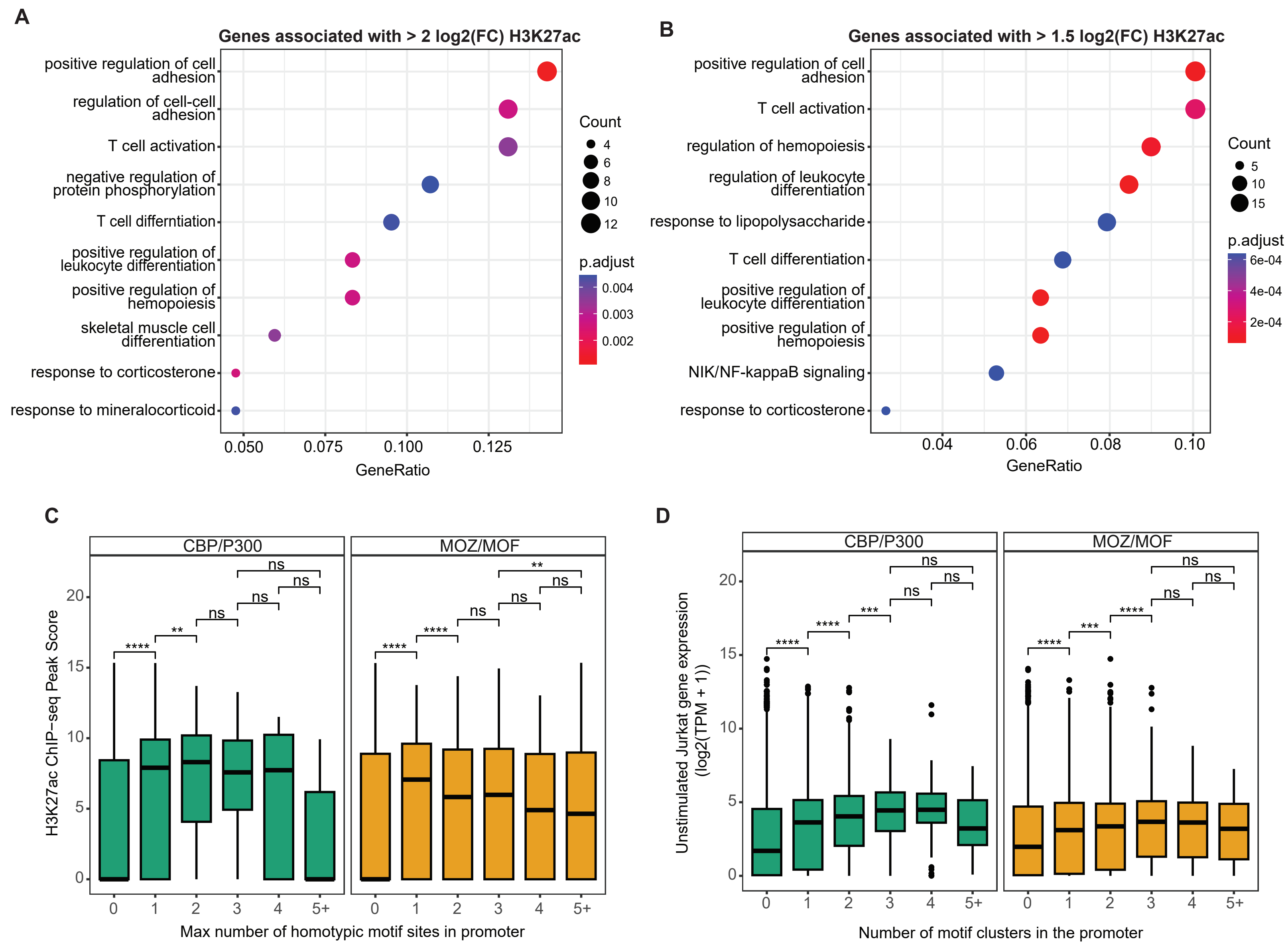
