## Supplemental Table 1 for "Rapid profiling of transcription factor-cofactor interaction networks reveals principles of epigenetic regulation"

| **Antibody** | **Company** | **Catalog #** |
| --- | --- | --- |
| **Primary Antibodies** | | |
| CBP | Abcam | ab10489 |
| P300 | Abcam | ab14984 |
| KAT2B | Santa Cruz | sc-13124 |
| GCN5 | Santa Cruz | sc-365321X |
| TIP60 | Santa Cruz | sc-166323 |
| MOF | Fortis/Bethyl | A300-992A |
| MOZ | Invitrogen | PA5-66742 |
| TBL1XR1 | Santa Cruz | sc-100908 |
| BRD4 | Fortis/Bethyl | A301-985A |
| **Secondary Antibodies** | | |
| Goat anti-mouse IgG (H+L) Highly Cross-Adsorbed Secondary Antibody, Alexa Fluor 488 | Invitrogen | A11029 |
| Goat anti-rabbit IgG (H+L) Highly Cross-Adsorbed Secondary Antibody, Alexa Fluor 647 | Invitrogen | A32733 |
